# Cholinergic regulation of sleep in the upside-down jellyfish *Cassiopea*

**DOI:** 10.1101/2024.10.04.616757

**Authors:** Michael J. Abrams, Aki Ohdera, Diana A. Francis, Owen Donayre, Henry Chen, Kevin Lu, Richard M. Harland

## Abstract

Perhaps nothing is stronger evidence of the importance of sleep than its conservation across animals [1], but the extent of its regulatory conservation is unknown. The upside-down jellyfish *Cassiopea xamachana* sleeps [2], and this behavior is controlled by radially-spaced marginal ganglia. After defining a sleep-wake threshold, we compared gene expression profiles of ganglia from animals sleep-deprived for two nights and found differential expression in many sleep-related genes including GABAergic, melatonergic, and cholinergic receptors. We focused on a nicotinic acetylcholine receptor alpha subunit-like (Chrnal-E), based on its differential expression, and selected animals for a second round of RNAseq that included both light-based and mechanically-based sleep-deprivation. Combining datasets revealed a short list of differentially expressed genes, of which *chrnal-E* is the most recognizable and well-supported, so we investigated its potential role in sleep regulation. First, we found that chemical cholinergic neuromodulators positively regulate pacemaker activity. Then, we showed by *in situ* hybridization that *chrnal-E* is expressed primarily within the ganglia, and that the area of expression expands after sleep deprivation. Next, we developed RNAi for use in *Cassiopea* and determined that Chrnal-E promotes wakefulness. Finally, we sampled circadian timepoints in the field and found in control conditions, *chrnal-E* has lowest expression late at night, but in sleep deprived animals, *chrnal-E* peaks at this time, supporting a link to wakefulness. Our finding that *Cassiopea* sleep is regulated by the cholinergic system underscores that mechanisms of sleep conservation are deeply conserved in animal evolution.

---

Sleep across animals is defined by three behavioral criteria: reversible quiescence, homeostasis, and latency-to-arousal [4]. This definition has mostly been applied to animals with centralized nervous systems; however, recently, sleep has been characterized in the jellyfish *Cassiopea* [2], and the polyp, *Hydra* [5], so sleep predates the emergence of centralized nervous systems, and likely serves a function in neural homeostasis. *Cassiopea* naturally orient upside-down on the shallow ocean floor (Figure 1B), and predominantly pulse in place through rhythmic contractions of the bell muscle to collect food, exchange gases, and support their symbiont [3]. The pulsing behavior is led by ganglionic pacemakers that reside in the rhopalia, which are radially-spaced light- and balance-sensing organs (Figure 1C). Cassiopea meet sleep criteria [2,4]: 1) They display quiescence, a slower pulse rate at night, that is rapidly reversible upon stimulation, 2) they respond slower to stimulation during sleep, evidence of increased latency-to-arousal, and 3) if deprived of this nighttime quiescence they experience a rebound, a compensatory low activity period, the following day (homeostasis). Because the ganglia control the pulsing behavior of the animal, the regulation of sleep must be integrated at these sites. Here we exploit *Cassiopea*, its distributed neural architecture, and its position as an early-branching metazoan, to investigate the emergence of sleep-regulatory mechanisms during evolution.

## Sleep-bout characterization in *Cassiopea*

To associate specific gene expression patterns with sleep or wake, we defined a behavioral activity threshold to separate the two states. Previously, *Cassiopea* were shown to increase latency-to-arousal at night, as measured by their response to gently falling from a stationary position in the water (drop-test) [2]. Here we find that low-activity (slow-pulsing) is associated with a reversible increased latency-to-arousal (i.e a sleep-like state). Using a similar approach to that applied in other animals [5-10]: 1) we determined a non-aroused activity level (*behavioral baseline*), and then 2) measured the response time to an arousing stimulus (Figure 1D). We take advantage of how *Cassiopea* briefly freeze pulsing after the drop-test, perhaps akin to the hyper-arousal freeze response of other animals [11].

We took the same pulse-counting approach as was previously published [2], and with modifications, monitored pulsation, quantifying the intra-pulse-interval (IPI) to calculate the average of 3-IPIs pre-dropping in both the light and the dark (Figure 1d2, Figure 1E); this established a non-aroused light and dark *behavioral baseline*, normalized by the mean activity. Then, for step 2, we measured the latency-to-arousal after the drop-test by quantifying how long it took the animal to begin pulsing (time to first pulse) after the drop (Figure 1E). After each 5 min acclimation and drop we repeated the test within 2 min to determine reversibility (Figure 1E). We found that slow pulsing animals (IPI>median), in both the dark and the light, had an increased latency-to-arousal for the initial test (Figure 1de2, right panel, blue), but not upon repetition (Figure 1e2, right panel, orange), indicating this test is identifying a reversible quiescent state. After 5 minutes animals acclimate, but during the 2 min post-drop arousal period, IPI is not indicative of their state, as both slow- and fast-pulsing animals respond quickly to the drop-test (Figure 1E). Together, these experiments show the slow steady-state pulsing activity in *Cassiopea* is indicative of a sleep state, so we moved on to investigate their regulatory response to nighttime sleep-deprivation.

**Figure 1.**
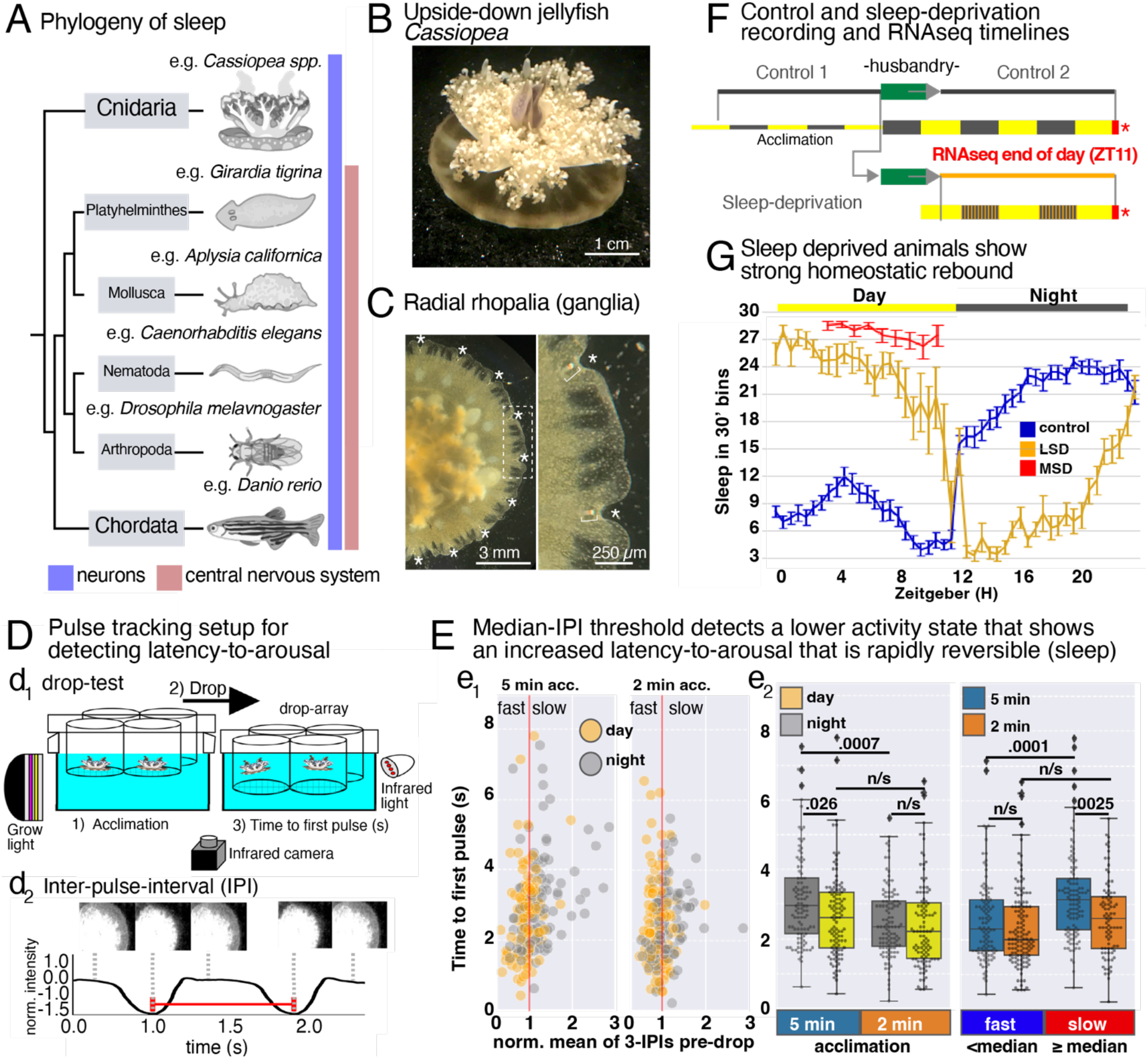
Phylogeny of sleep, *Cassiopea* pulse-tracking to detect latency-to-arousal and sleep bouts in *Cassiopea*. **(A)** Phylogenetic tree highlighting the six phyla with described sleep states, adapted from [16], and the lack of a CNS in *Cassiopea*. **(B)** *Cassiopea* in its natural upside-down orientation. **(C)** Oral view of *Cassiopea*, asterisks indicate the position of radially spaced rhopalia. **(D)** For drop-test recordings, **(d**_**1**_) jellies are placed in an array of clear screen-bottom cylinders and raised to a set height, then allowed to acclimate for 5 min. The array is then dropped, forcing the animals into the water column, which they eventually respond to by pulsing towards the bottom [2]. Recordings were made in infrared at 15 frames per second (CAM). **(d**_**2**_) The average pixel intensity is calculated every frame within a region of interest (example image series shown above the pulse trace) and normalized to remove background light fluctuations. The frames with the least intensity occur when the jelly is most contracted. An example time between contractions, the Inter-Pulse-Interval (IPI), is shown. **(E)** During the day (yellow) and the night (grey), on 8 jellyfish, across 4 days and nights, we cycled them through 8-10 rounds of 5 min for acclimation followed by a “drop,” measured their time to first pulse, and then quickly reset for a second drop to measure reversibility. The verticle red line is the normalized median of 3-IPIs pre-drop, the threshold between fast- and slow-pusling animals. **(e**_**1**_**) Left panel** is the result after 5 min of acclimation, and the **Right panel** is the result after repeating the experiment within 2 min. **(e**_**2**_**) Right panel**, box plots show the time to first pulse of fast-vs slow-pulsing animals, considering whether they are 5 min acclimated (blue) or a repeat experiment (orange). p-values calculated using one-way ANOVA, followed by Tukey HSD to adjust for multiple comparisons. **(F)** The behavioral recording paradigm is an initial recording (control 1), that includes 2 full nights and 2 full days, followed by a husbandry period for feeding and cleaning. Jellies are then returned to the recording setup for either 2 normal nights and 2 days of (control 2), or 2 nights of sleep deprivation (LSD). At the end of the recordings (ZT11, red bar and asterisk), animals have 3/4 of their ganglia amputated for RNAseq. For optimal RNA extraction, larger animals were used, ∼6cm diameter. **(G)** Average sleep in 30-min bins. control (blue), light-based sleep-deprived (LSD, orange), mechanically sleep deprived (MSD, red) error bars are SEM; control, n = 6 (288 hrs), LSD, n=7 (336 hrs), MSD, n=3 (72 hrs, no nighttime recording assessment, sleep-wake threshold set to pre-MSD median-IPI).

## Homeostatic response to sleep-deprivation in *Cassiopea* shows differential expression of genes similar to those in vertebrates

The molecular basis of sleep homeostasis is not well understood [12], nor is there much knowledge of its conservation across animals [1]. To target sleep homeostasis, we initially developed a light-based sleep-deprivation (LSD) method, giving the animals 5 min pulses of light every 25 min during the night. We carried out a series of behavioral recordings on six animals, three control and three LSD, and quantified the IPI (Figure 1d2, Figure F, Supplemental Figure 1A). We defined a sleep-wake threshold (norm. median IPI) from each 24-hour recording for each animal following the above method, giving us the amount of sleep in 30-minute bins. Control animals slept little during the day, except for a brief mid-day siesta, and sleep ∼70% of the night (Figure 1G, Supplemental Figure 1B). LSD animals showed a strong homeostatic rebound (compensatory low activity) during the day, and slept little at night, but eventually either habituated to the stimulus or were driven to sleep by the homeostatic system (Figure 1G, Supplemental Figure 1B).

We compared the gene expression profiles of animals in the late day (ZT11) that had either experienced control conditions or had been exposed to LSD for two consecutive nights, to attempt to measure a regulatory response to sleep deprivation and detect sleep-related genes (Supplemental Figure 1B). Several core neuroactive receptors, and many components of GABAergic, circadian, and cholinergic pathways were represented in the transcriptome (Supplemental Figure 2A-D). Principal component analysis clustered control animals together with one LSD animal, and the two other LSD animals clustered apart, suggesting variation in individual response to LSD (Supplemental Figure 1d1).

**Figure 2.**
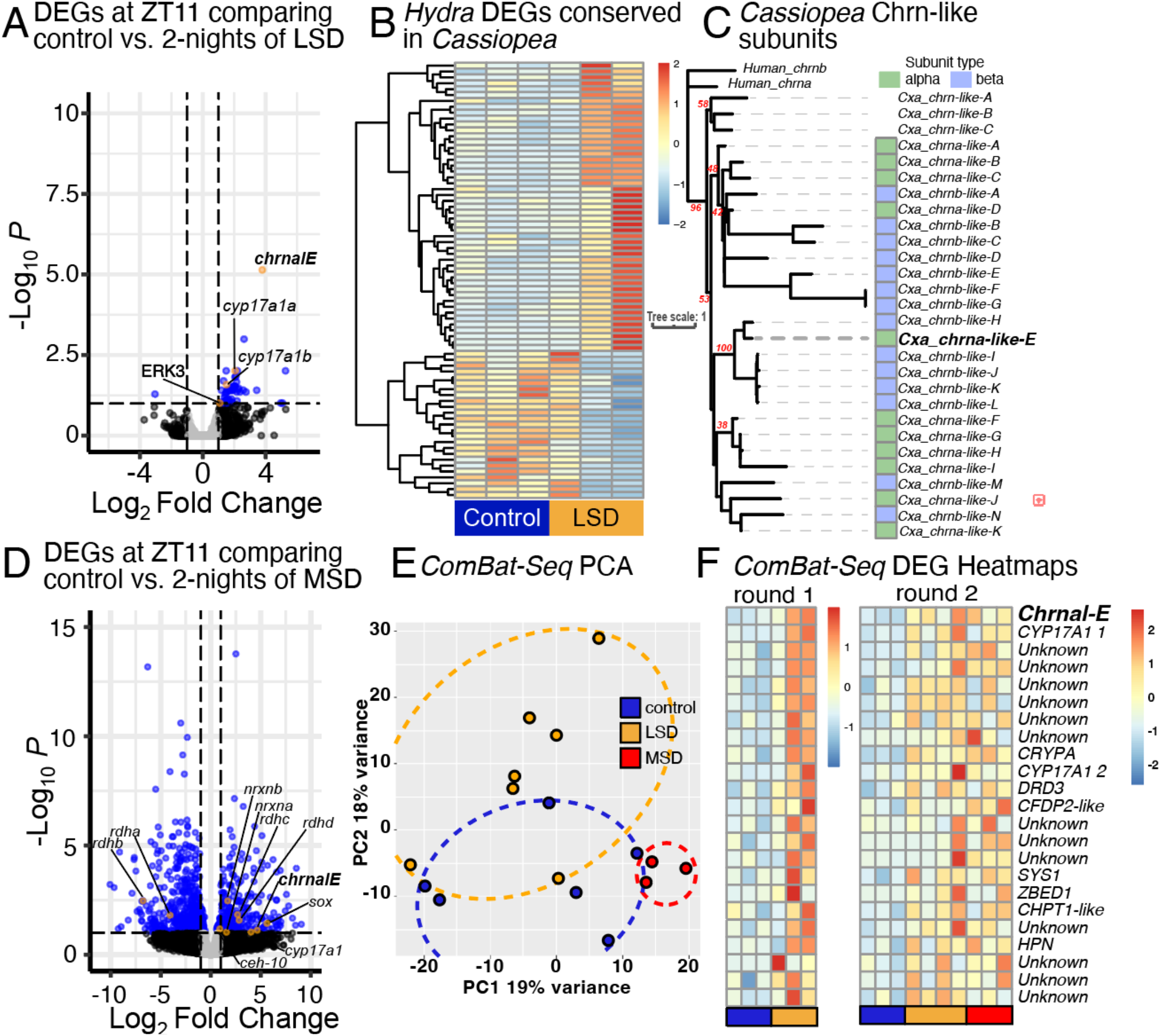
RNAseq reveals sleep-related DEGs and a potential. **(A)** Volcano plot visualizing log_2_ fold-change (x-axis) and log_10_ p-adjusted value (y-axis) of gene expression from a pairwise comparison between light sleep deprivation (LSD) treated jellyfish and control jellyfish. Dotted lines show a p-adjusted cutoff of 0.1 and a fold-change cutoff of (+/-) 1. Genes meeting these thresholds are shown with a blue dot, with key genes highlighted in orange with labels. **(B)** Heatmap of regularized log transfromed gene expression of sleep-related genes conserved between *Hydra* and *Cassiopea* [4]. **(C)** Phylogenetic analysis of known and putative Chrns. Maximum likelihood phylogeny generated in IQ tree. Bootstrap support in red. **(D)** Volcano plot visualizing gene expression from a pairwise comparison between mechanical sleep deprivation (MSD) treated jellyfish and control jellyfish. Dotted lines show a p-adjusted cutoff of 0.1 and a fold-change cutoff of (+/-) 1. Genes meeting these thresholds are shown with a blue dot, with key genes highlighted in orange with labels. **(E)** Principle component analysis of ComBat-Seq normalized expression data for meta-RNAseq analysis. Samples largely clustered according to their treatment. Colored ellipses were manually applied to highlight clusters. RNAseq samples in blue are controls, orange are LSD animals, and red are MSD. **(F)** Heatmap of regularized log transformed gene expression for the twenty-four differentially expressed genes determined with DEseq2 using the ComBat-Seq normalized reads analysis in the LSD vs control (round 1) dataset and the MSD vs control (round 2) dataset. Genes are listed according to the p-adjusted values calculated with DESeq2.

Nevertheless, DEG analysis identified 62 up-regulated genes and 1 down-regulated gene (Figure 2A) (Supplemental Table 1). Using sleep-associated *Hydra* genes [5] to prioritize *Cassiopea* orthologs, we found clear differences between LSD samples and controls, although one LSD treated sample more closely resembled control patterns of gene expression (Figure 2B, Supplemental figure 1d1). Differentially expressed genes included those associated with sleep deprivation from other organisms, including ERK3, steroid-metabolizing enzymes, and a neuronal acetylcholine receptor (Figure 2A). The neuronal acetylcholine receptor was one of the most up-regulated genes in response to LSD. We also performed KEGG enrichment analysis and found seventeen enriched pathways, including sleep-related pathways “cholinergic synapse” and “NOD-like receptor signaling pathway”, as well as immune response pathways “TNF signaling pathway”, “NF-kappa B signaling pathway”, and “IL-17 signaling pathway” (Supplemental Table 2).

Given the general importance of cholinergic systems in animal neural function and sleep homeostasis [13-16], we chose to explore the gene further. *Cassiopea* has 25 orthologs of acetylcholine receptor (Figure 2C), which can be categorized by the presence of a well characterized Cys-Cys pair ligand binding site, as well as the presence of a C-loop [17]. We phylogenetically identified 11 alpha subunits (we named chrnal-A-K) and 14 beta subunits (we named chrnbl-A-N) of cholinergic receptors in the *Cassiopea* genome, named so as not to confuse them with the established subunits (Figure 2C). The differentially expressed ortholog (*chrnal-E*) was identified as an alpha subunit.

## Mechanical sleep deprivation also induces chrnal-E expression

To determine whether mechanical stimuli result in gene expression responses in *Cassiopea* to light stimuli, we repeated the RNAseq experiment using mechanical sleep deprivation (MSD). After optimizing flow, containment, and imaging for this condition, we found animals were mechanically sleep deprived using pulses of water, 1 minute every 4 minutes, during the night, which drove significant rebound sleep during the day (Figure 1G, Supplemental Figure 1B). Given the variable gene expression response to light-based sleep deprivation, we sleep deprived more animals, and selected animals based on their *chrnal-E* expression (Supplemental Figure 1e1). MSD animals exhibited a greater number of differentially expressed genes (353 up-regulated, 600 down-regulated; p-adjusted = 0.1) (Figure 2D, Supplemental Table 3). As with the first experiment, individual response to MSD varied, despite careful sample collection (Supplemental Figure 1e2). As expected, *chrnal-E* was up-regulated (4-fold Log2) in sleep deprived animals (Figure 2D), as well as a *neurexin*, a second acetylcholine receptor (chrnbl-J), steroid-metabolizing enzymes, a series of development genes (*homeobox, cranio-facial development protein, sox*), and genes involved in retinoid metabolism (*retinol dehydrogenases*). KEGG pathway enrichment analysis identified pathways involved in neural regulation (“Axon guidance”, “Axon regeneration”, and “Synaptic vesicle cycle”) among genes that were down-regulated (Suppmental Table 4)

An alternative method to compare datasets is Combat-Seq, which adjusts raw RNAseq counts by modeling batch effects between multiple datasets using a negative binomial regression [18]. We assessed both Round 1 and Round 2, which included a second set of LSD animals. Correcting for batch effect with *ComBat-Seq* led to samples from both control, LSD, and MSD animals clustering closely within a PCA space (Figure 2E). Using conservative cutoffs, we recovered 24 genes that showed consistent expression differences between control and sleep deprived animals.

*Chrnal-E* emerged as the top-most differentially expressed gene. Among the list of genes were steroid alpha-hydroxylase/lyase-like, craniofacial development protein, as well as a dopamine receptor (Supplemental Table 5). Although many of the genes were not differentially expressed when the two experiments were analyzed separately, the 24 genes from Round 1 and Round 2 reveal clear patterns of up-regulation in SD animals (Figure 2F). These data suggest *Cassiopea* sleep homeostasis is likely regulated by *chrnal-E* amongst other genes.

## Conserved receptor structure and response to cholinergic modulators identifies a bonafide nicotinic cholinergic receptor in *Cassiopea*

Cnidarian Chrna7-like and 10-like showed strong similarity in BLAST results for *Cassiopea chrnal-E*, which show striking structural and domain similarity with Human Chrna7 (Figure 3A). Further, the structure gives insight into potential functional similarity, as the charged residues in the extracellular vestibule that regulate calcium permeation in Human Chrna7 [19] are also present in only this *Cassiopea* Chrna subunit (Figure 3a1&a2).

**Figure 3.**
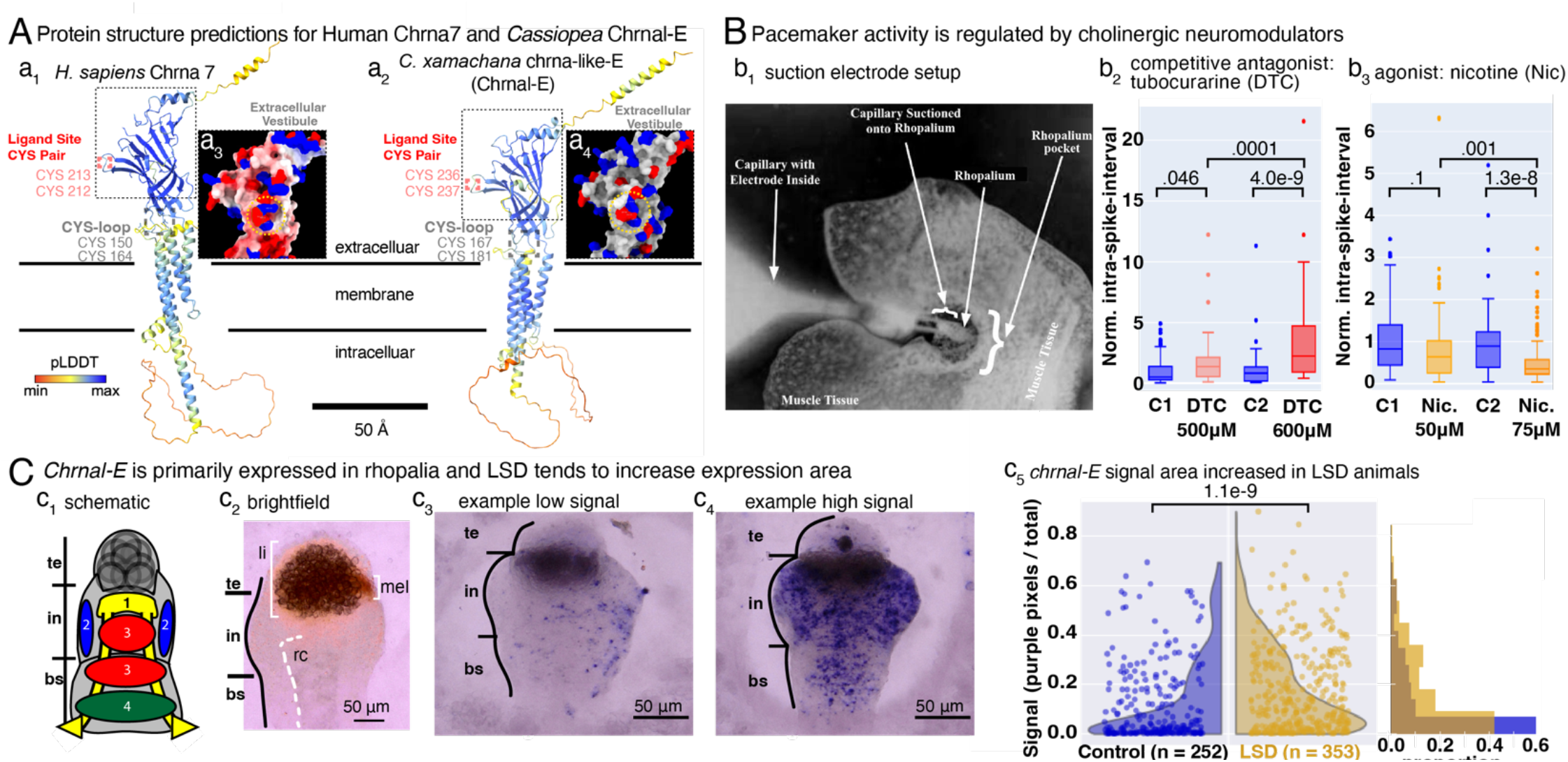
Evidence of an intact cholinergic regulation in the nervous system of C*assiopea*. **(A)** Alpha fold protein structure predictions for **(a**_**1**_) Human Chrna-7, and **(a**_**2**_) *C. xamachana* chrma-like-E (Chrnal-E), with labeled ligand site and CYS-loop indicating alpha-subunit identity. Extracellular, membrane, and intracellular regions demarcated. pLDDT color legend indicates structure confidence. The charged residues in the extracellular vestibule that regulate calcium permeation **[11]** for **(a**_**3**_) Human Chrna-7, and **(a**_**4**_) Chrnal-E. Color is charge, red (negative), and blue (positive). **(B)** Electrophysiology: amputated rhopalia were suctioned into glass capillary electrodes (**b**_**1**_) and action potentials (APs) were recorded, and the inter-spike interval (ISI) was calculated. Explants recorded in artificial seawater (control, blue) versus artificial seawater with dissolved **(b**_**2**_) tubocurarine at 500 μM or 600 μM (salmon), a competitive antagonist against nicotinic acetylcholine receptors or dissolved nicotine at **(b**_**3**_) 50 μM or 75 μM (orange), an antagonist for nicotinic acetylcholine receptors. p-value calculated using One-way ANOVA, with Tukey’s HSD applied post-hoc. **(C)** *chrnal-E* expression shifts towards greater ganglionic expression when sleep deprived. **(c**_**1**_) Schematic of schyphozoan rhopalium ganglionic neural groups adapted from *Aurelia* [24]. te, terminal, in, intermediate, and bs, basal segments. **(c**_**2**_) Brightfield image of a rhopalium, oral view, with the three labeled segments. The left half of the radial canal, rc, is marked by a white dotted line, which brings circulation to the rhopalium. The dark brown lithocyst, li, is in the terminal segment, and the light brown melanin, mel, of the spot ocelli on the aboral side is situated between the intermediate segment and terminal segment. Expression pattern of *chrna*, with examples of low **(c**_**3**_), and high-**(c**_**4**_) expression, and their respective signal quantification. **(c**_**5**_) Quantification of ganglionic staining; ganglia are selected from images, and the ratio of purple pixels to total pixels in the ganglia is found. **Left panel**: violin plot of the distribution; **Right panel**: normalized histogram. control, n = 252, SD, n=353. p-value calculated using Kolmogorov-Smirnov test.

The cholinergic system is critical to sleep regulation in worms, flies, fish, mice, and humans [13-16]. Human Chrna7 is associated with memory and Alzheimer’s disease, which are both strongly linked to sleep [20]. Interestingly, one Chrna is required for *Drosophila* sleep and promotes sleep in response to an increasing homeostatic drive (the increasing need for sleep during prolonged wake) [13]. Recently, Chrns were found to have neural and non-neural roles in the cnidarian anemone *Nematostella* [21]. Additionally, cholinergic pace-maker neurons in a freshwater cnidarian *Hydra* were characterized [21] as similar to the pacemaker neurons in the vertebrate gut [22]. However, the cholinergic system has yet to be implicated in sleep regulation of prebilaterian lineages [5], so we were driven to investigate the role of Chrnal-E in regulating sleep in *Cassiopea*.

While cholinergic receptors are present, they may not control neurological activity, so we used suction electrode electrophysiology on amputated ganglia (Figure 3b1, Supplemental Figure 1G-H) to measure the time between action potentials, the intra-spike-interval, in the presence of nicotinic acetylcholine receptor effectors. We bath loaded tubocurarine (DTC), a competitive antagonist, or nicotine (Nic), a highly membrane permeable agonist. In a dose-dependent manner, tubocurarine increased the intra-spike-interval (Figure 3b2; 500 μM, p-value .046; 600 μM p-value, 4.0e-9, DTC 500 μM < DTC 600 μM, p-value .0001) while nicotine decreased the intra-spike-interval (Figure 3b3; 50 μM, p-value .1; 75 μM p-value, 1.3e-8, Nic. 50 μM > Nic. 75 μM, p-value .001). These chemicals act on nicotinic acetylcholine receptors at the site of the ligand binding, so we consider these data strong evidence of acetylcholine acting as a neurotransmitter in *Cassiopea*.

## *Chrnal-E* is primarily expressed in ganglia and transcript localization area increased due to LSD

We also confirmed that *chrnal-E* is expressed in the rhopalia. Rhopalia are divided into three segments [24]: terminal, intermediate, and basal (Figure 3c1). The terminal segment includes the lithocyst for balance, and the indentation between the terminal and intermediate segment contains the pigmented-spot ocelli on the aboral side, with its brown melanin pigment. The basal segment connects the rhopalium to the body of the animal. *Chrnal-E* localizes to the oral side of ganglia (Figure 3c3-4), predominantly in the intermediate and basal segments, but without a singular discrete pattern. There is considerable variability in mRNA expression patterns, both in levels and in ganglionic regions, even between the ganglia of a single animal (Supplemental Figure 3D). Subcellular localization of RNA appears non-nuclear and asymmetrical within the cytosol (Supplemental Figure 3E).

Using *in situ* hybridization allowed us to detect the location of *chrnal-E* expression, and though the intensity of the color is not necessarily indicative of expression level, we did strive to understand if LSD caused a difference in *chrnal-E* pattern. We selected the rhopalium from each *in situ* image, pooled the images, and measured the ratio of the purple pixels to the total number of pixels (signal), in each rhopalium (Supplemental Figure 3C). We found a significant shift towards broader *chrnal-E* expression in LSD ganglia (Figure 3C, p-value 1.1e-9). Because of its association with ganglia, and its change in response to LSD, we were motivated to ask if Chrnal-E is functionally involved in regulating sleep.

## RNAi reveals a functional role for Chrnal-E in promoting wake in *Cassiopea*

We optimized RNAi conditions based on previous works [25-27] (Supplemental Figure 4A-G), and then tested whether the *chrnal-E* knockdown affects sleep behavior (Supplemental Figure 4H, I). We inserted 387bp of *chrnal-E* into the L4440 vector. As a control, we used 427bp of *mcol*, which we had previously characterized both using *in situ* hybridization (Supplemental Figure 3b3,4), and which we considered unlikely to affect sleep behavior. For characterization of the knockdown, five animals were fed the *chrnal-E* vector, and five animals were fed the empty vector, for two weeks. Staining of the *chrnal-E* knockdown rhopalia reveled reduction in expression area of the *chrnal-E* mRNA (Figure 4A, p-value 3.3e-6).

**Figure 4.**
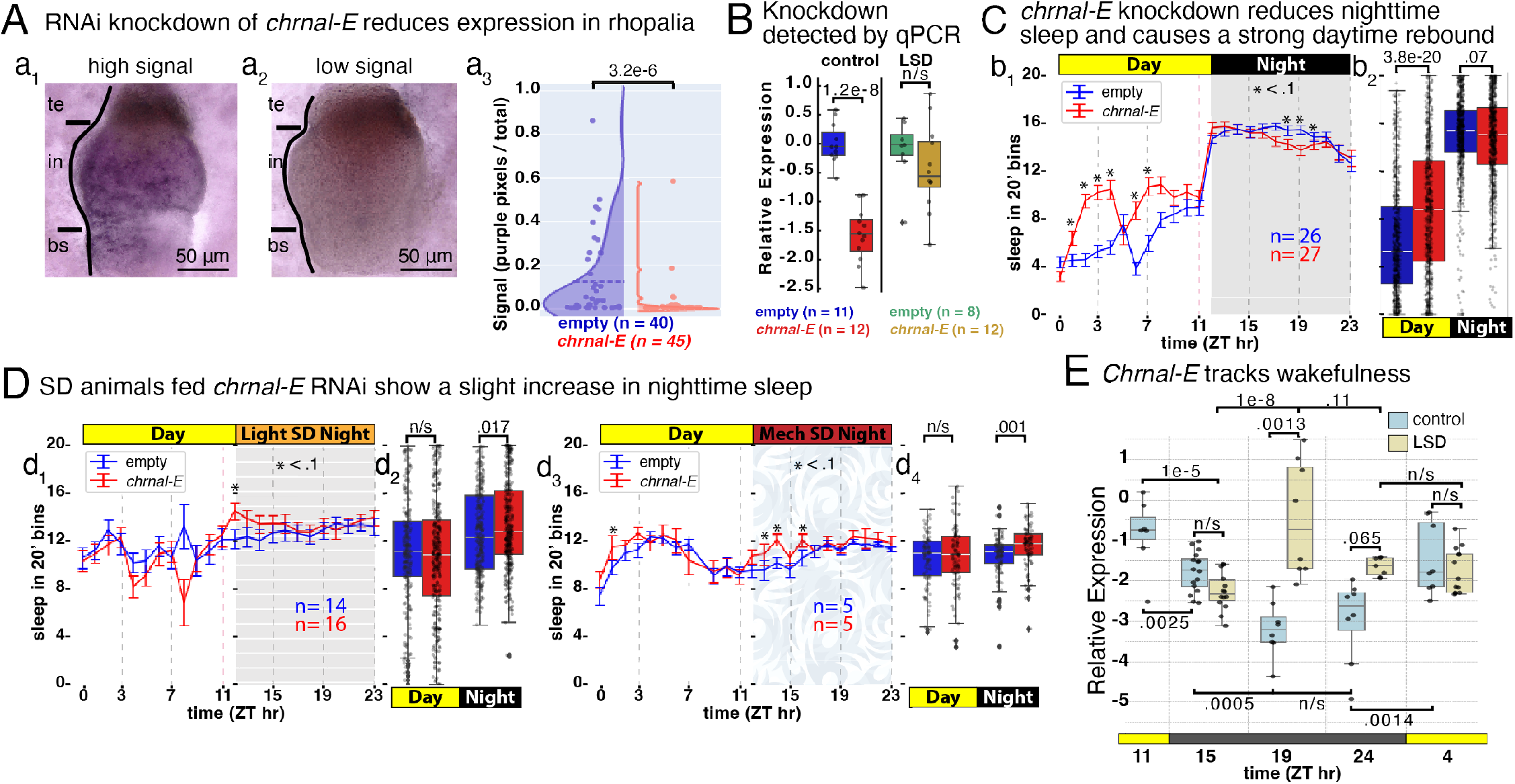
RNAi knockdown *chrnal-E* increases daytime sleep. **(A)** *In situ* hybridization characterization of *chrnal-E* in animals fed RNAi empty-vector (control, 5 animals) or *chrnal-E* (5 animals) for two weeks; **(a**_**1**_) high and **(a**_**2**_) low, *chrnal-E* expression. **(a**_**3**_) Purple-pixel quantification of ganglionic staining. p-value calculated by One-way ANOVA. **(B**,**D)** Sleep in 20’ bins, showing the average sleep of animals fed empty-vector (blue) or *chrnal-E* (red), for control **(b**_**1**,**2**_), LSD **(d**_**1**,**2**_), MSD **(d**_**3**,**4**_). Sleep per hour and **(b**_**1**_, **d**_**1**_, **d**_**3**_) sleep day vs. night **(b**_**2**_, **d**_**2**_, **d**_**4**_**). (E)** Detection of circadian expression of *chrnal-E* comparing controls to LSD animals from field samples. Error bars are SEMs. p-values generated by One-way ANOVA, and Tukey’s HSD post-hoc test was applied.

Next, we recorded pulsing behavior for 3 weeks, during which the animals were fed RNAi vectors, and finally, at ZT 11, tissue was extracted and the expression level of *chrnal-E* quantified. We detected a strong reduction in *chrnal-E* expression in animals fed *chrnal-E*-RNAi (Supplemental Figure 4G). Behaviorally, by the second week, animals fed *chrnal-E* vector began displaying increased daytime sleep and decreased nighttime sleep compared to *mcol*and empty-vector RNAi (Supplemental Figure 4H,I). After 3 weeks, we found that *chrnal-E-*vector fed animals strongly increased their daytime sleep (Figure 4B). Animals that were sleep deprived, either using LSD or MSD, did show sleep deprivation or daytime rebound compared to controls (Figure 4d2,d4). When we quantified *chrnal-E* expression in LSD animals at ZT11, we did not find a significant knockdown (Figure 4C); however, animals fed *chrnal-E* RNAi vector did show a slight increase in nighttime sleep compared to controls (Figure 4D). These findings indicate Chrnal-E drives wakefulness because knock-down animals slept more than they typically would, both during the day as in control conditions, and at night during sleep deprivation.

## *Chrnal-E* expression tracks wakefulness

The RNAi result showed a strong daytime effect wake, leading to increased sleep; coupled with the previously noted variability in *chrnal-E* expression detected at ZT11, we hypothesized that *chrnal-E* expression is dynamic. To address this, we sampled many *Cassiopea*, in the field at the Key Largo Marine Research Laboratory, at circadian timepoints starting at the end of the day (ZT11) through to the morning (ZT4) and measured *chrnal-E* expression using qPCR (Figure 4E). We found *chrnal-E* in control animals decreased at ZT19 and stayed low until it increased again in the morning (ZT4) (Figure 4E), indicating a possible connection to the circadian cycle, or it could be part of the homeostat response, as *chrnal-E* expression is higher during wake (day) and lower at night (sleep). Strikingly, in the middle of the night (ZT19), *chrnal-E* expression is much higher in animals that experienced 7 hours of LSD (Figure 4E, p-value .0013). So, rather than staying at a daytime level, which we may expect if it were light driven, or decreasing as it does in control, *chrnal-E* drastically increased expression as the animals experienced prolonged wake. Perhaps, as the animals began to habituate or sleep through the light stimulus as they do in the lab (Figure 1G), *chrnal-E* levels reduced by the end of night (ZT24), but it was still elevated compared to the control (Figure 4E, p-value .065). This indicates *chrnal-E* directly promotes wake as part of the sleep homeostat: *chrnal-E* increased with sleep-debt but was lower during frequent sleep.

Finally, the circadian expression informs the behavior of the RNAi knockdown animals in control and sleep deprived conditions. The knockdown reduced the daytime level of *chrnal-E*, at ZT11, by approximately 1.5-fold (Figure 4C, p-value 1.2e-8) compared to controls, which is similar to the 3-fold reduction we detected in the circadian timepoints ZT19-24 (Figure 4E, p-value 1e-9). Second, while the LSD animals did not show a significant expression knockdown at ZT11 (Figure 4C), perhaps the expression at ZT15-19 would be reduced enough to suppress *chrnal-E* upregulation and reduce the nighttime wakefulness of the sleep deprivation. Together, these findings indicate that *chrnal-E* promotes wake and is a key member of the homeostatic sleep system in *Cassiopea*.

## Conserved regulation of sleep in *Cassiopea*

The molecular mechanism of sleep homeostasis remains a mystery and a subject of intense research in the sleep field [28,29]. We endeavored to uncover components of the sleep homeostat in *Cassiopea*. We found that sleep deprived animals tend to change expression of a similar set of genes related to sleep across animals, providing support for their being an ancestral sleep-regulatory system. We discovered a nicotinic acetylcholine receptor alpha subunit, *chrnal-E*, that is highly differentially expressed in association with sleep deprivation. RNAi knockdown revealed that chrnal-E drives wakefulness, as control animals slept more during the day, and LSD and MSD animals slept more at night during sleep deprivation, when they would normally be deprived. Finally, tracking its expression level from day to night, we found that *chrnal-E* in control animals has lowest expression at ZT19, while LSD animals peak at that time, indicating that *chrnal-E* increases expression in line with wakefulness. Therefore, we hypothesize that *chrnal-E* acts in the sleep homeostat by tracking and promoting wake.

We recognize, however, that the differential expression observed does not appear in every animal, or every ganglion (Figure 2). While there may be technical explanations, cnidarians also exhibit variability in response to environmental stimuli at the individual animal level for metamorphosis [30], self-repair [31], regeneration [32] and the behavioral response to sleep deprivation [2] (Figure 1G, Supplemental Figure 1B). While the prevalent changes provide insight into sleep regulation, the animals that don’t respond as anticipated may yet be compensating in an unknown way.

Phylogenetic analysis of cnidarian Chrns showed they grouped together, separate from those in Bilateria, and the addition of *Cassiopea* subunits did not particularly alter previous phylogenetic structure [21], supporting the notion that Chrns independently radiated in cnidarian and bilaterian lineages. However, there is some debate as to whether the presence of Chrns is evidence of a true cholinergic system. Our readout for their behavioral state is their pulsing activity, which is controlled by pacemaker cells in the ganglia, and because *chrnal-E* mRNA is also present there, we hypothesized a neural regulatory function for *chrnal-E*. Indeed, tracking pacemaker generated action potentials (APs) in the presence of cholinergic modulators supports our finding, as tubocurarine decreases AP frequency and nicotine increases AP frequency, a competitive antagonist and agonist, respectively. Together, these findings join other works [21,33] in supporting the presence of an intact cholinergic system in Cnidaria, with a role for acetylcholine as a neurotransmitter.

To know where *chrnal-E* exists in the sleep regulatory pathway will require more investigation. Chrnal-E could track sleep debt as a homeostat in normal conditions, similar to Redeye [13]; or it could interact with the circadian clock [29]. There is also evidence for cell-autonomous sleep-regulatory mechanisms of a non-neuronal nature in mammals and *Drosophila* [1]: tissue metabolism, growth or oxidative stress could drive sleep, so perhaps *chrnal-E* acts downstream of a stress sensor. We are excited by how RNAi opens the door to targeted further studies that should resolve sleep-regulatory mechanisms and the extent to which there is conservation in *Cassiopea*. To date, there are numerous examples of conserved regulation of sleep, and with evidence indicating some conserved functions of sleep, our findings support there being a conserved form of sleep in the Cnidarian-Bilaterian ancestor. Further, because *Cassiopea* sleep is cholinergic, it emphasizes the deep importance of this system in the evolution of sleep and wake regulatory mechanisms.

## Author Contributions

M.J.A. conceived of the project. M.J.A. and R.M.H oversaw the project. M.J.A., A.O., D.F., O.D., H.C., K.L., conducted experiments. M.J.A., carried out behavioral analysis. M.J.A and A.O. performed the gene expression analysis. M.J.A., D.F., H.C developed and executed, and O.D., performed, molecular biology protocols. M.J.A conceptualized, designed, and built experimental setups. M.J.A and K.L developed and performed electrophysiology experiments. M.J.A wrote the paper with input from A.O., D.F., and R.M.H.

## Acknowledgment

We thank Lea Goentoro from Caltech, CA; and Victoria Sharp from Pennsylvania State University, PA, for generously supplying *Cassiopea* medusa and polyps. We thank David Raizen for critical reading of the manuscript. We thank Marta Truchado Garcia and our Harland Lab colleagues for their counsel and technical expertise. We thank our compatriots in the International Cassiopea Workshop, CassiopeaBase, and the KLMRL, for input and discussion. We thank the incredible UC Berkeley undergrads, without whom there would be no project, who worked on *Cassiopea* molecular biology, Casey Herbert, Katie Cheng, Alexander Garcia-Calvillo, Brian Benito, Kamron Emami, Rithvik Ghankot, Braeden Lemm, Stella Frank, and Alisson Kennedy; and *Cassiopea* behavior, Hannah Zeigler, Celeste Hsu, Tessa Auer, Sofia Telfer, Jerry Yu, Matthew Lim, Mayra Arellano, Shuangyue Li, and Mira Patel; and *Cassiopea* electrophysiology, Lilian Zhang, Brandon Lee, William Ling, and Hailey Mozier. Thank you to the UC Berkeley URAP system. Thank you to Nicole King for initial advice and support. This work was supported by the Miller Institute at University of California, Berkeley, and the C.H. Li Distinguished Professor fund.

## Materials and Methods

### Pulse Tracker

#### Recording setup and approach

Light/Dark tests and RNAi behavioral experiments used 4in x 4in x 4in acrylic boxes filled with enough sea-water to cover the jellyfish (approximately 1.5 inch of water height) with 28-32 ppt artificial seawater. Four square watch glasses of 1 and ⅝ inch (Carolina Biological Supply) were placed in each box, with each watch glass holding one jellyfish for a maximum of four jellyfish per container. White netting was placed above and to the sides of the jellyfish to prevent mixing animals during the recording, and to ensure that the jellyfish remained within their watch glass. Acrylic boxes were placed in a Pyrex dish filled with distilled water and with three Pulaco PL-188 heaters set to 84°F. This system resulted in a heated water bath for the jellyfish. The lights used for the behavioral recording jellyfish were fixtures with two full spectrum T5 and two actinic T5 bulbs. The lights were on a 12 hour cycle from light to dark, where lights came on at 7 AM (ZT0) and turned off at 7 PM (ZT12). SD experiments utilize bright LED lamps attached to a cycling timer that turns on 5min every 25min. The same camera and lens was used, but the Matlab program recorded mp4 videos at 15fps.

To analyze pulse rate of jellyfish, recorded videos were processed with the program FFmpeg [34], using custom parallelized version, ffmpeg_parallel.sh. This converts multiple videos into image stacks simultaneously at a rate of 15 fps. Pulse Tracker Analysis Software [2] can bused used on any length recording period, for control vs SD RNAseq experiments we analyzed the full recording and used larger animals with ∼6 cm diameter. For RNAi experiments there were more animals, and they had ∼3 cm diameter. For RNAi, we analyzed 20 minutes of every hour, equivalent to 18000 images, throughout the experiment as was done previously [2]. There are two main steps in the Pulse Tracker Analysis Software. First, Pixel Intensity Extractor takes the first image of the first time period and allows the user to select a rectangular region of interest surrounding the jellyfish. Then, inside of this region, the software measures average pixel intensity of each frame of that period. Second, Peak Finder, normalizes the output from step one, and then detects local minima that correspond with muscle contractions. When the intensity passes a chosen threshold value a pulse is demarcated, indicated with a red dot (Supplemental Figure 1a2,3). The output to this program has the time at which a contraction occurred and the IPI between the current and previous pulse. Ideal Peak Finder (Supplemental Figure 1a2) vs problematic Peak Finder (Supplemental Figure 1a3), where the average pixel intensity change was not significant enough to allow for clear pulse detection. The low-quality graphs can occur due to insufficient pixel change difference when the jellyfish pulsed, which often was caused by lighting fluctuations surrounding the jellyfish independent of pulse behavior, or the animal managed to turn sideways. We did not have issues with the larger animals in the RNAseq experiment but did have some issues with the smaller animals for RNAi. We attempted to choose 20-minute periods with good pulse quality and dropped time points from individuals if no satisfactory 20 min period could be found.

### Confocal Imaging

Images were taken with a Zeiss LSM 880 with confocal microscope at 10x, 20x, and 40x magnification, or using a Zeiss LSM 700. Image processing (Z projection, color display) and quantifications were done with Fiji software.

### *In situ* hybridization

#### Probe Synthesis

Plasmids of our genes of interests were extracted from colonies stored in our bacterial glycerol stock library. 3-5 ug of plasmid DNA were digested using select restriction enzymes. An absence of additional cut sites, which could be present within the insert or in another section of the vector, was verified prior to choosing the appropriate digestion enzyme; NcoI, NotI, or SphI were used with SP6 polymerase while SpeI was used for T7 polymerase. Following a 3-3.5 hour 37°C digestion, the corresponding polymerase was used to synthesize DIG-labeled RNA probes.

#### In situ hybridization

Our rhopalia specimens were stained by *in situ* hybridization using a protocol adapted from *Nematostella* [35]. Briefly, animals were anesthetized in 800 μM menthol. Animals were first fixed in 4% PFA, 0.02% Tween-20, 0.3% glutaraldehyde, in PBS, pH 7.2. Fixation times vary by probe. For *rfamide, chrna*, and *mitf* we used 30 min, *minicollagen* required 40 min. Fixations were gently rocked at room temperature. After the first fixation, specimens were washed in PBS, and then put through an ethanol series to permeabilize and de-pigment the specimens. Samples were placed in 50%, up to 100% ethanol, in 4 stages, and back down, with 5 minutes per stage, with gentle rocking. Then, a second fixation was used, this time in 4% PFA in PBS, pH 7.2, bringing the total fixation time to 1 hour per probe. Specimens were washed several times, and then treated with Proteinase K (50 μg/mL) in 0.1% Tween-20 PBS (PTw), for 23 minutes. After stopping the reaction with glycine, specimens were washed then treated with 1% triethanolamine (TEA), 1% TEA + 3 μ/mL acetic anhydride, 1% TEA + 6 μ/mL acetic anhydride, and then washed in PTw. Then specimens are refixed in 4% PFA for 1 hour, and then washed in PTw. Samples were then transferred to hybridization buffer, first washing at room temperature, then brought to 65°C, for 24 hours, for pre-hybridization. Probes (at 1μg/mL) were denatured at 80-90°C for 5 min, and after removal of hyb. buffer from samples, probes were added and the samples returned to 65°C for 48 hours. After retrieving probes, which can be re-used indefinitely, samples were washed in descending concentrations of hyb. buffer mixed with 2x SSC pH 7.0, down to 25% hyb. buffer and 75% 2X SSC, in 30 minutes steps, and returning samples to 65°C. After 100% 2X SSC washes, samples were washed in descending concentrations of 0.02X SSC down to 25% 0.02X SSC, 75% PTw, and finally 100% PTw, initially at 65°C, then at room temperature. Samples were then transferred to Roche Blocking Buffer, and placed on a rocker at 4°C overnight. Anti-Digoxigenin was diluted in Roche Blocking Buffer 1:5000, and returned to 4°C overnight. After several washes in 1% Triton-X100 in PBS, samples were transferred to AP buffer with MgCl_2_. Then samples were transferred to AP buffer without MgCl_2_, and then finally into the color reaction, 3.3 μL/mL NBT (50mg/mL) and 3.3 μL/mL BCIP (50mg/mL), or Fastred. Reactions were kept in the dark, but monitored, and generally took 3-5 hours at room temperature to complete. Samples were then washed in PTw and fixed for 1 hour in 4% PFA in PBS at room temperature and stored in PBS at 4°C for several months without issue.

#### *In situ* hybridization controls

*Cassiopea* rfamide mRNA staining was consistent with the antibody staining with localization to the ganglia and sensory nerve net [36], in medusae, and in the tentacles and mouth of the polyp (Supplemental Figure 3B1&2). In a contrasting control, nematocyst-specific gene *minicollagen* (*mcol*) mRNA stain showed the classic salt-and-pepper pattern in the oral side of the bell, and the tentacles and body of the polyp (Supplemental Figure 3B3&4), similar to *Nematostella* [37].

#### Imaging

Specimens were mounted on glass slides, or in agarose-mold dishes, and imaged on a Keyence VHX-5000 digital microscope.

### Immunohistochemistry

Medusae or amputated ganglia were fixed in 4% paraformaldehyde for 2 hours at room temperature (20-25 °C), and washed in PBS. To remove pigment from *Symbiodinium microadriaticum*, a dinoflagellate algal symbiont of Cassiopea, samples were dehydrated and rehydrated in an ethanol series (50%, 70%, 90%, 100%, 100%, 90%, 70%, 50%) for 5-minute intervals followed by a final wash in PBS. For samples stained with 488-Phalloidin, isopropanol substituted for ethanol for 30 second intervals. Samples were blocked in PBS containing 10% CAS-Block for 2 hours. Primary antibodies FMRFamide (EMD Millipore AB15348; rabbit, 1:1500), or acetylated Tubulin (Sigma-Aldrich MFCD00164512; mouse, 1:200), were diluted in 10% CAS-Block, and samples were incubated at 4 °C with gentle rotation overnight. Sample was washed with PBS, then placed in secondary antibody mixtures, depending on the primary antibody, Alexa Fluor 488 (mouse, 1:200), Alexa Fluor 555 (rabbit, 1:200), dilutes in 10% CAS-Block, and incubated in the dark at 4 °C for 12 hours. Sample was washed PBS, and nuclei stained using 1ug/ml Hoechst 33258 (ThermoFisher 62249) for 2 hours.

### RNA extraction and quantification

#### RNA extraction

For RNAseq, 3/4 of the rhopalia from each animal (approximately 12 per animal) were pooled for each sequencing sample. For qPCR, 6 rhopalia were amputated, and 2 rhopalia were pooled per animal in 3 separate extraction. After amputation, specimens were transferred into Stellar Scientific Deadbolt 1.5 mL tubes, and were immediately flash frozen in liquid nitrogen and stored in a -80°C freezer. For extraction, tubes were seated on a liquid-nitrogen-cooled mortar and were homogenized physically with a cooled pestle (Thomas Scientific). The Plant Spectrum Total RNA kit (Sigma) was used to isolate the RNA from the homogenized specimens. Ground tissues were immediately vortexed in 1 mL of Lysis buffer, double the amount specified by the kit. After material from the binding solution was applied to the columns, they were treated with a DNase digestion (Sigma). When needed, samples were additionally cleaned and concentrated using the Zymo Research RNA Clean and Concentration Kit and eluted into 13 uL. Once concentrated, samples of RNA were analyzed with an Agilent Bioanalyzer system to assess yield and integrity.

### RNA sequencing and differential gene analysis

Paired-end 150 bp RNA sequencing was performed at the UC Berkeley sequencing core facility using the Illumina NovaSeq 6000. Reads were trimmed and quality filtered using Trimmomatic (version 0.39) with default settings. Trimmed reads were aligned to the gene model predictions of the *Cassiopea* genome version 0.2 [38], accessed through the Joint Genome Institute genome resource portal (**https://my-cocosm.jgi.doe.gov/Casxa1/Casxa1.home.html**) using STAR (version 2.5.3a) with the following settings: -- limitOutSJcollapsed 1000000 --limitSjdbInsertNsj 1000000 --outFilterMultimapNmax 100 --outFilterMismatchNmax 33 --outFilterMismatchNoverLmax 0.3 --seedSearchStartLmax 12 --alignSJoverhangMin 15 -- alignEndsType Local --outFilterMatchNminOverLread 0 --outFilterScoreMinOverLread 0.3 --winAnchorMultimapNmax 50 --alignSJDBoverhangMin 3. Differential expression analysis was performed using DESeq2 [39]. The Wald test was used for hypothesis testing, with FDR cut-off of 0.05. Visualization of the differential expression analysis was performed with regularized log transformed data. To identify a universal sleep deprivation response in gene expression of LSD and MSD animals, raw reads from experiment 1 and experiment 2 were batch corrected using ComBat-Seq function in the Bioconductor package sva version 3.42.0 [18]. Batch corrected counts were used as input for differential expression analysis using DEseq2 as described above.

### Gene identification

Primarily using *Nematostella* and *Clytia*, genes of interest were BLAST against the *Cassiopea* transcriptome, and hits then reciprocally BLAST for secondary confirmation. Domain analysis was performed using HMMER [40]. Phylogenetic analysis was performed using phylemon2 [41], and trees were constructed using maximum likelihood analysis (MLA) from protein sequences.

### Phylogenetic analysis

38 putative nAChR genes were predicted in the Cassiopea genome. A maximum-likelihood protein alignment of putative nAChRs was performed with MAFFT (version 7.429) and manually filtered following previously described criteria [21]. Sequences were checked for the C-loop domain, with cysteine residues separated by 13 amino acid residues. Sequences lacking the motif were excluded from further analysis. Sequencing with over 75% of the alignment absent from the TM domain were also excluded. After filtering, 24 protein coding sequences were retained. Sequences with a cysteine-cysteine pair prior to the transmembrane domains were classified as alpha subunits. Phylogenetic reconstruction of putative Cassiopea AChR was performed with iqtree (v2.0.3) with 2000 ultrafast bootstrap replications.

Protein sequences alignment of nicotinic acetylcholine receptor subunits were performed with MAFFT (version 7.429) with default settings. Sequences were retrieved from NCBI, except for *Nemopilema nomurai* [42] and *Clytia hemisphaerica* [43], which are available through the respective data repositories. Downloaded sequences shorter than 390 bp were removed prior to alignment. Alignments were trimmed using BMGE (version 1.2) with default settings. Reconstruction of a maximum likelihood phylogenetic tree was performed using IQ-TREE (version 2.2) [44], for which model selection was performed with ModelFinder [45]. Phylogenetic reconstruction was performed using ultrafast bootstrap approximation with 2000 replicates.

### Cloning

#### cDNA Synthesis

cDNA was prepared using Superscript IV Reverse transcriptase and RNA samples of at least 100 ng.

#### qPCR Primer validation and analysis

Experiments were performed with a Bio-Rad CFX Real-Time Machine. We used cDNA samples made from >200ng of RNA to assess primer efficiency. Serial cDNA dilutions of 40%, 20%, 10%, 5%, 2.5%, and 1.25% were prepared. cDNA and primers for the genes of interest were added to reactions with SsoAdvanced Universal SYBR Green supermixes and were prepared in duplicates for Standard Curve and Relative Quantification experiments. After validation, for every sample, *bactin, btubulin, mcol*, and *chrnal-E* were amplified. A minimum of three animals were used in each comparison, with three biological samples taken from each animal, and three technical replicates were used in every qPCR run. qPCR results were analyzed following standard qPCR calculations and error propagation [46].

### RNAi procedure in Cassiopea

#### RNAi vector assembly and induction

The RNAi construct synthesis and induction is based on techniques employed in the planarian *Schmidtea mediterranea* [25], and the nematode *C. elegans* [26], the freshwater polyp *Hydra Vulgaris* [27]. As shown in Supplemental Figure 4, to generate RNAi constructs, sequences (∼300-900 bp) from the genes of interest were inserted between the T7 promoters of the ampicillin-resistant, double T7 L4440 vector, and these recombinant vectors were transformed into HT115(DE3) *E. coli* (tetracycline-resistant, rnc14::ΔTn10, containing lacUV5 promoter-T7 polymerase). An overnight culture of L4440 in HT115(DE3) (grown in LB with 100 ug/mL ampicillin and 12.5 ug/mL tetracycline) was diluted 1:100 in 2xYT (100 ug/mL ampicillin) and incubated at 37°C until the OD_600nm_ = 0.4. To induce the culture, isopropyl β-D-1-thiogalactopyranoside (IPTG) was added to a final concentration of 1mM before a 5-hour incubation at 37°C. 2 ml of the culture was pelleted, the supernatant removed, and then stored at -80°C.

#### RNAi cocktail prep

Decapsulated brine shrimp eggs (Brine Shrimp Direct) were pipetted into a 1.5 mL tubes, and using a tabletop centrifuge briefly spun down, and the excess water was removed leaving approximately 300 uL of brine shrimp eggs. Eggs were crushed with a pestle using an electric tissue homogenizer, or a ceramic mortar and pestle, until the eggs no longer made a cracking sound. Following complete emulsification, we added 1 uL drop of blue food coloring, and 50uL of 2% PEG with dissolved blue chalk, in order to visualize the food inside of the jellyfish. We then added 400 uL of 0.8% low melting point agarose dissolved in sterile filtered sea water (SFSW) to the emulsified eggs. We transferred this egg mixture to its respective bacterial pellet; bacteria pellets are set to 125 uL (if pellet is too large or small it is adjusted). We then transferred this to a respective extrusion syringe, which correlated with each different treatment, with brine shrimp netting securely fastened to the tip. The filled apparatus was left to solidify on ice. We rapidly compressed the syringe to extrude the mixture into a 1.5 mL tube, then the mixture was diluted 1:1 in SFSW. RNAi cocktail was stored in 4 °C for up to 1 week.

#### RNAi feeding schedule

Every day at approximately 1PM (Zeitgeber hour 6) the jellyfish received 50-100 uL of the RNAi cocktail. If it was on a recording day, we removed the jellyfish from the recording setup prior to feeding and placed them in their respective weekday cubbies in a husbandry setup, and if not on a recording day we fed them directly in their weekday cubbies. We used a P200 pipette to slowly extrude the RNAi cocktail onto the oral arms of the jellyfish, and to verify eating we could visualize the blue food inside of the mouths of the jellyfish. The jellyfish were allowed 1 hour to eat, and then the excess food and mucus was removed. If recording, animals were placed back in the recording setup, and if on a non-recording day, they were kept in the husbandry set up.

## Supplemental Figures

**Supplemental Figure 1.**
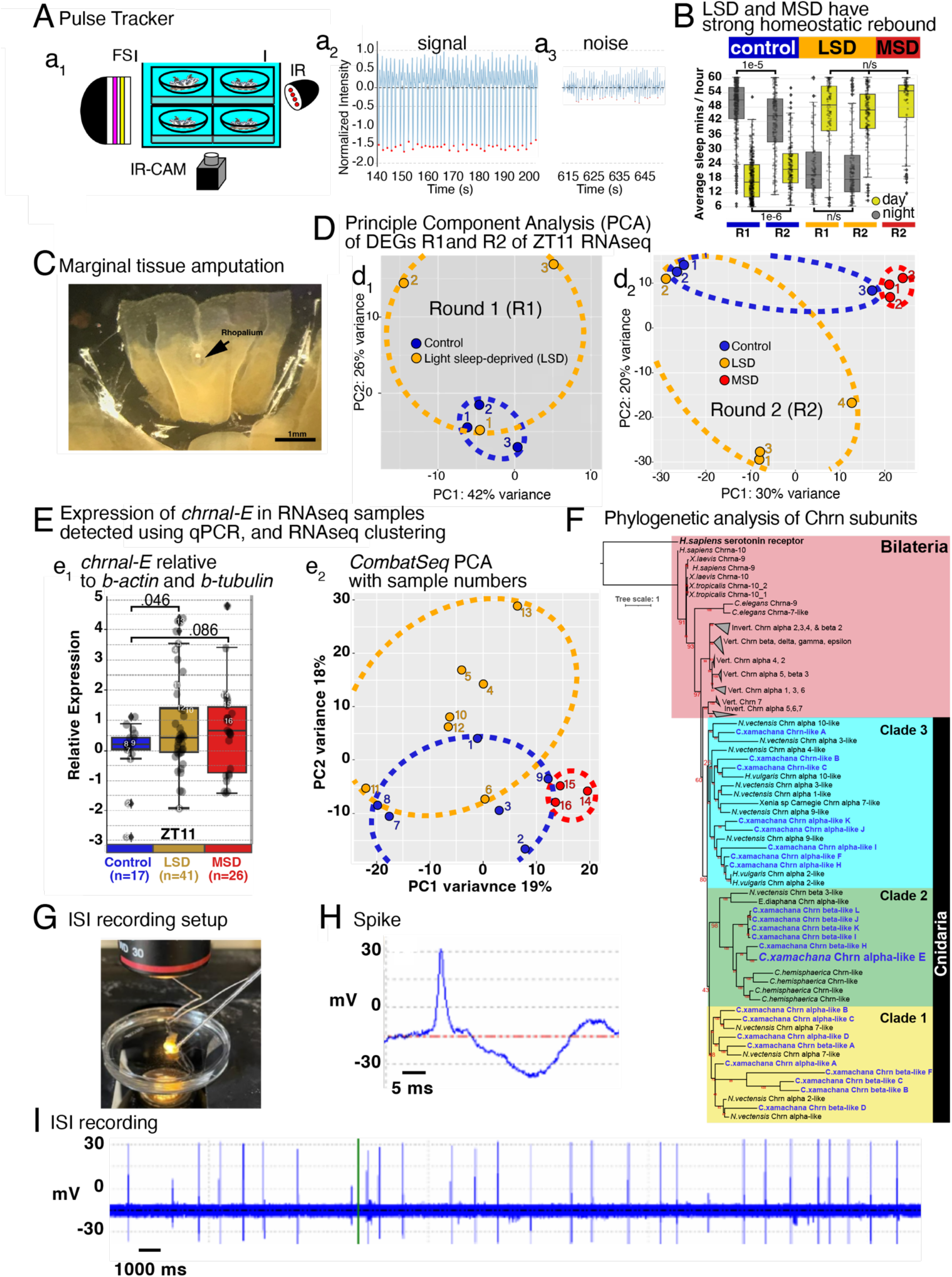
Pulse Tracker recording and analysis, rhopalia amputation, and RNA quality control. **(A)** Pulse Tracker recording setup, **(a**_**1**_) with full spectrum (FS) and infrared (IR) laterally angled lights and camera recording at 15 frames per second, fitted with an IR filter. **(a**_**2**_) Code then finds the troughs demarcating contractions, and labels them with red dots if the peak is over a set threshold, in this case -0.5 of the normalized intensity, and exports files with the time of each contraction and the inter-pulse-interval (IPI). **(a**_**3**_) Example of unusable pulse tracker data where there is not sufficient change in average pixel to clearly label contractions compared to noise from background. **(C)** Amputation of rhopalia and surround tissue, while avoiding thicker mesoglea-rich tissue. **(D)** Principle component analysis of RNAseq, samples plotted in PCA space. **(E)** qPCR to confirm differential expression of *chrnal-E* at ZT11 in control, LSD, and MSD conditions. Numbers samples in **(e**_**1**_) are the samples represted on the PCA in **(e**_**2**_). Control (blue), LSD (orange) and MSD (red). p-values generated by One-way ANOVA. **(F)** Phylogenetic analysis of known and putative Chrns. Maximum likelihood phylogeny generated in IQ tree. Human seratonin receptor is used as out-group for our analysis. *C. xamachana* sequences are in blue text on tips of tree branches.**(G)** Reference wire and electrode suctioned onto rhopalium in a petri dish with temperature regulated sterile filtered seawater on a light microscope state. **(H)** Depolarizing exgraceulluar waveform identified as spike. **(I)** Electrophysiology recording.

**Supplemental Figure 2.**
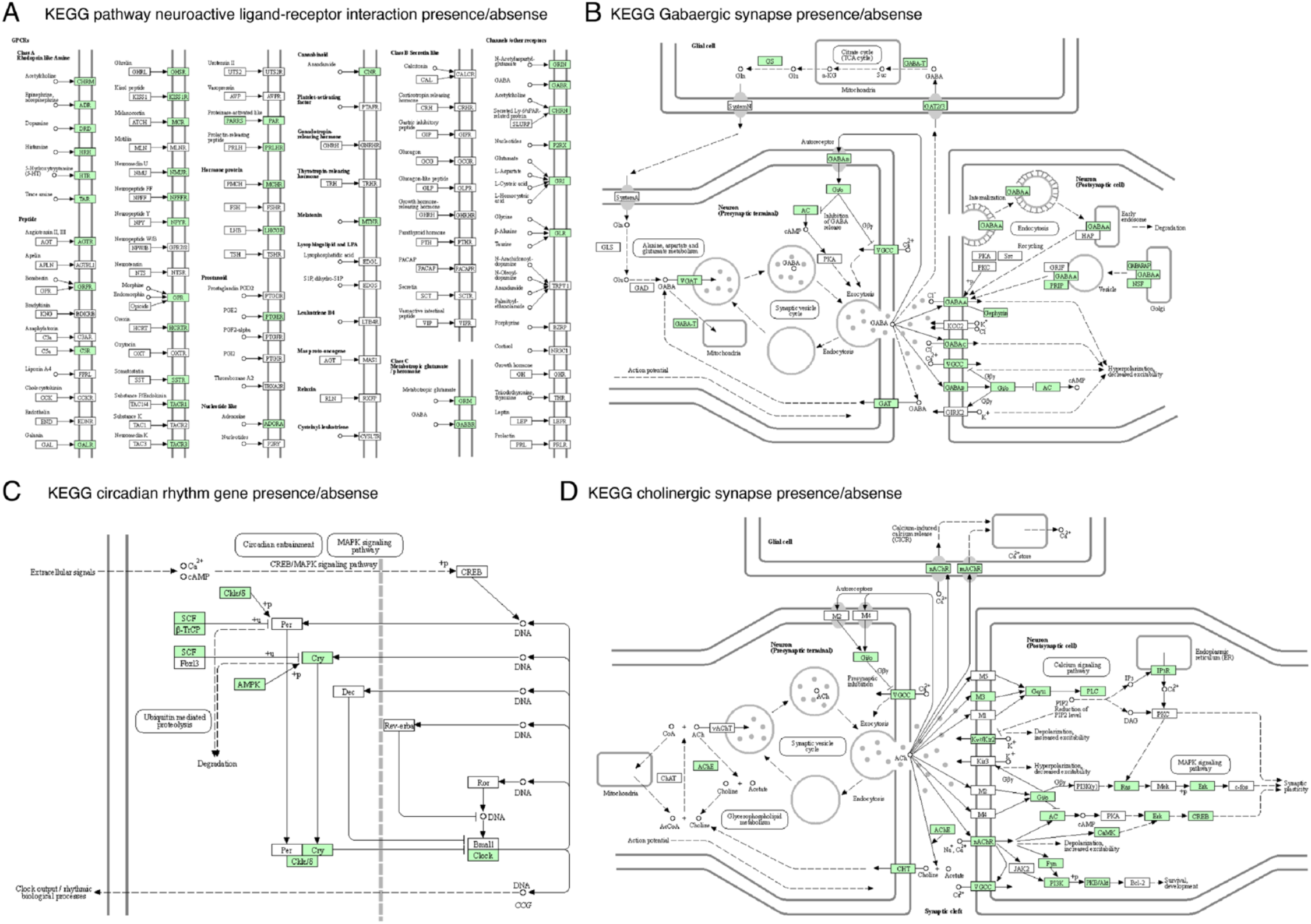
KEGG pathway presence, absence. **(A-D)** KEGG pathways showing presence (tan-box)/absence (white-box) of **(A)** neuroactive ligand-receptor interactions, **(B)** GABAergic synapse, **(C)** circadian rhythm, and **(D)** cholinergic synapse.

**Supplemental Figure 3.**
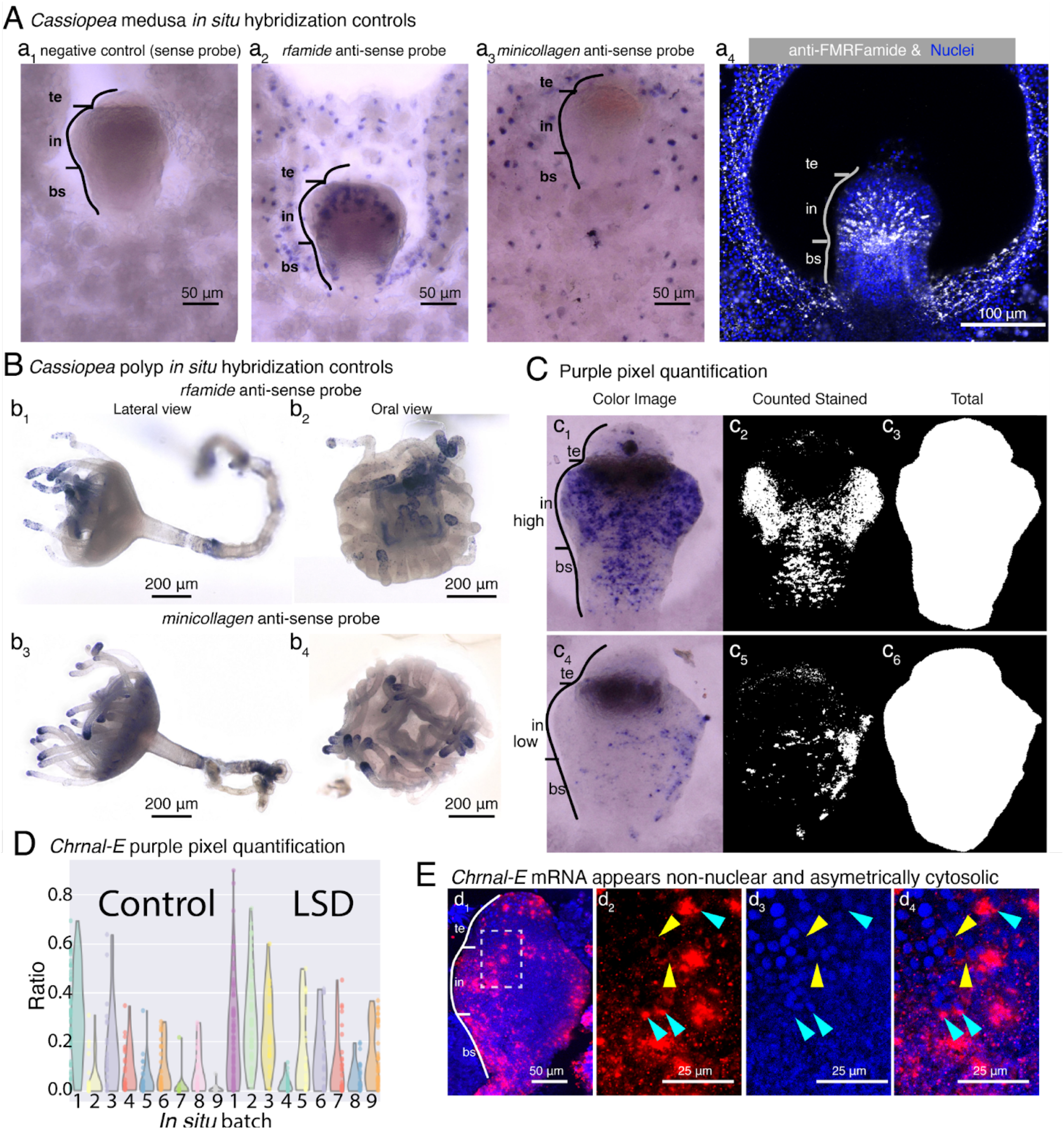
*in situ* hybridization of *rfamide* and *minicollagen* in *Cassiopea* polyp and rhopalia, *chrnal-E* expression quantification and localization. **(A)** *in situ* validation in *Cassiopea*. **(a**_**1**_) *Cassiopea* rhopalia show very low background staining with sense probes. **(a**_**2**_) *rfamide* is highly expressed within the rhopalia and in the pocket that surrounds it, consistent with sensory nerve localization [24,25]. **(a**_**3**_) *Minicollagen (mcol)* is not expressed in the rhoplalia, instead it has a salt-and-pepper pattern consistent with nematocyst localization. **(a**_**4**_) FMRFamide antibody labels neurons in the diffuse nerve net and the ganglia, with a similar pattern to the *rfamide* anti-sense probe, and previous reports [24,25]. **(B)** *rfamide* and *mcol* expression patterns in polyps. **(b**_**1**_) *rfamide*, and **(b**_**2**_) *mcol*, localizing to the tentacles and around the mouth. However, *rfamide* is more dispersed in the tentacle and is less expressed in the body than *mcol*. **(C)** Purple pixel quantification of *chrnal-E* expression. **(c**_**1**,**4**_) Color images of chromogenic *in situ* hybridization. **(c**_**2**,**5**_) A mask for blue-purple pixels is generated. **(c**_**3**,**6**_) A mask for all pixels within the rhopalium is generated. Signal is the ratio of blue-purple pixels in rhopalium/ total pixels within the rhopalium. **(c**_**7**_) Quantification of staining per ganglion per jellyfish, and the aggregate per condition in the box plot. p-value calculated Wilcoxon rank-sum. **(D)** Fastred *in situ* hybridization of *chrnal-E* to visualize subcellular localization. **(d**_**1**_) Fastred *chrnal-E* expression pattern on the oral side of a rhopalium. **(d**_**2**_) Fastred *chrnal-E*, **(d**_**3**_) Hoeschst nuclear stain, **(d**_**4**_) merged. Expression of *chrnal-E* appears non-nuclear, and asymmetrically cytosolic. Yellow arrows indicate low levels and cyan arrows show high levels.

**Supplemental Figure 4.**
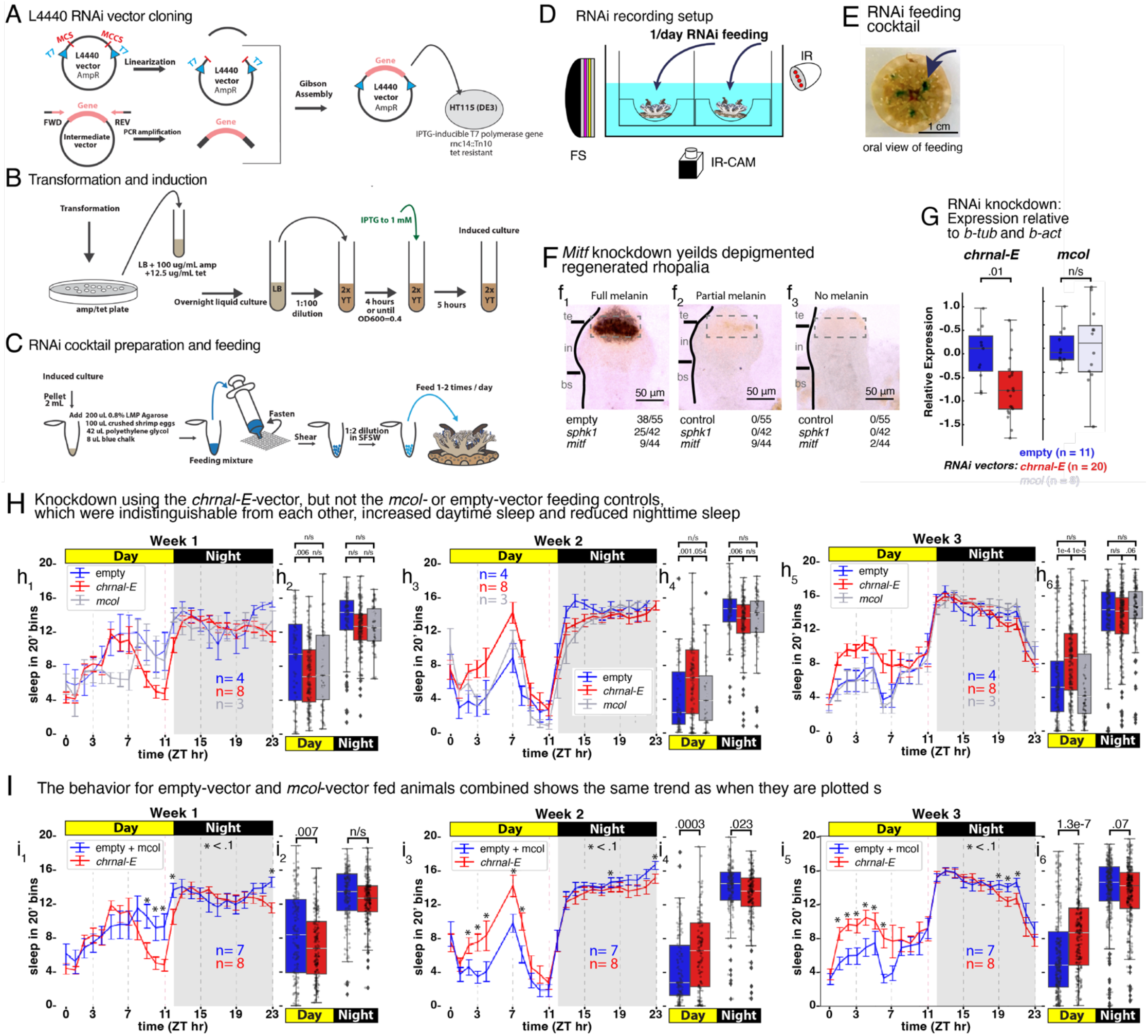
RNAi vector cloning, induction, and preparation, and RNAi behavioral recording setup and group and RNAi knockdown of *mitf* and *chrnal-E*. **(A)** ∼250-1000bp of target gene are amplified from cDNA and cloned into the L4440 vector, transformed into high efficiency DH5alpha, sequenced, then re-transformed into HT115 *E*.*coli*. **(B)** Induction using 1 mM IPTG in 2xYT media is accomplished in a series of grow-out steps. **(C)** Induced cultures are pelleted and stored at -80C. When needed, pellets are thawed, and mixed with 0.8% LMP, crushed brine shrimp eggs, PEG, and chalk, and extruded through a brine shrimp filter, and then diluted in sterile filtered artificial seawater. RNAi food can be kept at 4C for up to 1 week, feedings are once or twice a day, depending on the size of the animal and its tolerance of the cocktail. **(D)** RNAi can be fed to animals in the Pulse-Tracker recording setup. **(E)** RNAi-food is placed directly on the tentacles, once or twice per day, which have oral groves that bring the food to their manubrium. **(F)** The RNAi was fed daily to three animals each condition for 21 days. All ganglia were removed after one week. After two weeks of recovery regeneration and melanin pigmentation was screened. There is a significant reduction in regeneration in animals fed *mitf* compared to empty (chi-square, p-value .017). Of the regenerated ganglia **(f**_**1**_) only *mitf* animals had two additional phenotypes, **(f**_**2**_) partial melanin, and **(f**_**3**_) no melanin. **(G)** qPCR of *mcol* and *chrnal-E*, relative to *beta-actin* and beta*-tubulin*, comparing expression in animals fed empty vector versus those fed *mcol* or *chrnal-E* vector. p-value calculated by One-way ANOVA. **(H)** % sleep, day and night, for week1-3. Animals fed empty-vector (blue), mCOL-vector (orange), and ACHRa-vector (grey). Empty- and mCOL-vectors do not show significant differences in sleep at any time point in week 2 or week 3. Error bars are SEM. **(I)** Behavior from empty- and mcol-vectors are combined to generate a combined control behavior, to compare against the *chrnal-E* knockdown.

**Supplemental Figure 5.**
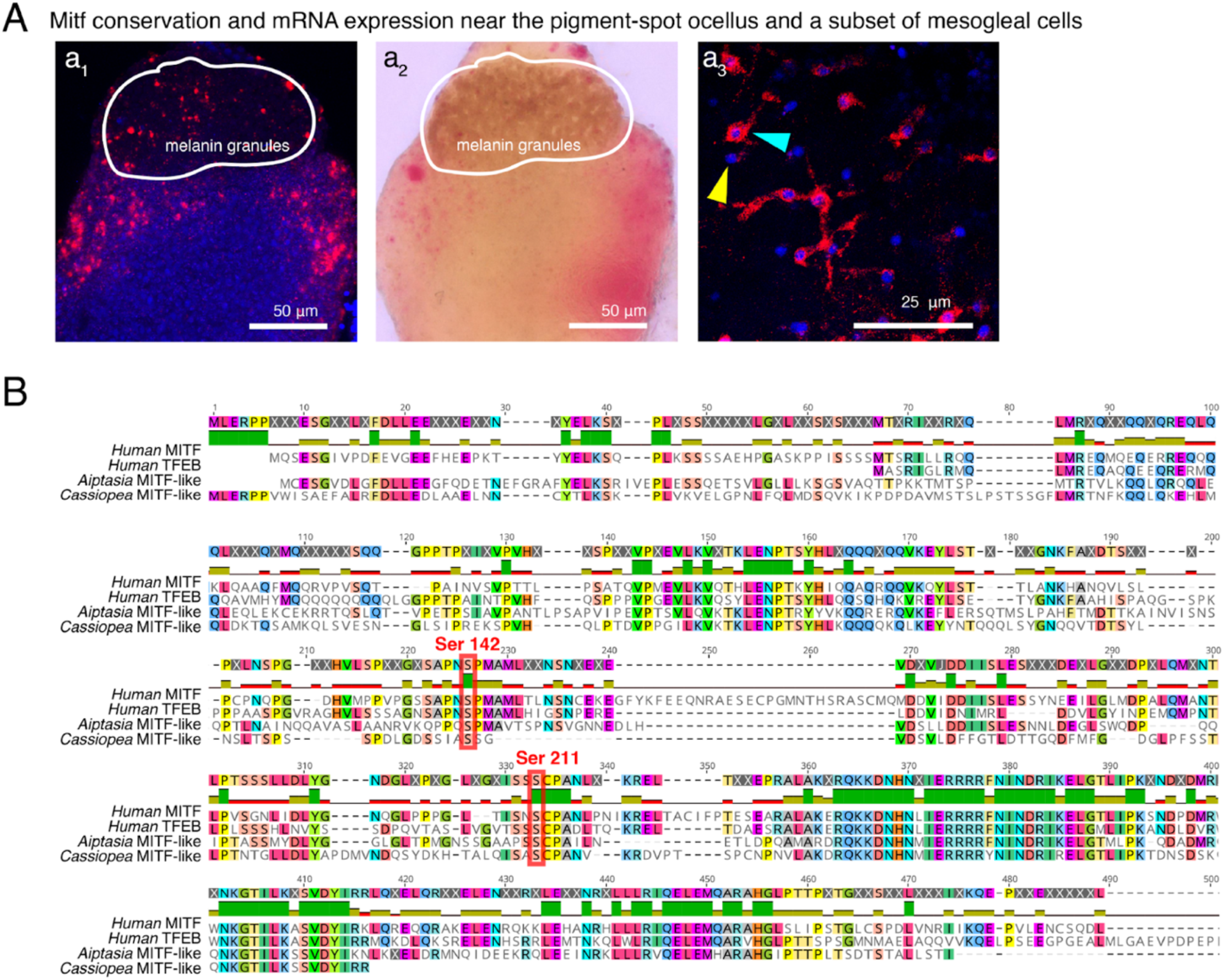
*Mitf* expression localization and sequence conservation. **(A)** *Mitf* expression in *Cassiopea*. **(a**_**1**_) *Mitf* mRNA (red), is expressed flanking, and internal to, the region of melanin granules of the pigmented-spot ocellus **(a**_**2**_**). (a**_**3**_) *mitf* mRNA also localizes to the cytoplasm of some cells inside the mesoglea, at high (cyan) and low (yellow) levels, compared to other mesogleal cells that appear just a nuclei (blue). **(B)** *Mitf* shares conservation of mTOR phosphorylation sites, Ser142 and Ser211, red boxes) and is therefore likely to be controlled by mTOR, as is the case with Human *mitf* [47,48].

